# Historic and contemporary museum specimens implicate Northern Red-backed Vole (*Clethrionomys rutilus*) as borealpox host as early as 1990s

**DOI:** 10.64898/2026.03.22.713527

**Authors:** Maya Juman, Jeffrey B. Doty, Clint N. Morgan, Audrey Matheny, Ariel Caudle, Marissa Breslin, Natalie M. Hamilton, Aren Gunderson, Katherine Newell, Julia Rogers, Victoria A. Balta, Italo B. Zecca, Florence Whitehill, Faisal S. Minhaj, Molly M. McDonough, Adam Ferguson, Yu Li, Crystal Gigante, Yoshinori Nakazawa, Joseph McLaughlin, Link E. Olson

**Affiliations:** Department of Veterinary Medicine, University of Cambridge, UK; Department of Mammalogy, University of Alaska Museum, University of Alaska Fairbanks, Fairbanks, AK, USA; Poxvirus and Rabies Branch, Division of High-Consequence Pathogens and Pathology, National Center for Emerging and Zoonotic Infectious Diseases, Centers for Disease Control and Prevention, Atlanta, GA, USA; Alaska Division of Public Health, Section of Epidemiology, Anchorage, AK, USA; Epidemic Intelligence Service, Centers for Disease Control and Prevention, Atlanta, Georgia, USA; Arctic Investigations Program, Division of Infectious Disease Readiness and Innovation, National Center for Emerging and Zoonotic Infectious Diseases, Centers for Disease Control and Prevention, Anchorage, AK, USA; Department of Biological Sciences, Chicago State University, Chicago, IL, USA; Gantz Family Collections Center, Field Museum of Natural History, Chicago, IL, USA

**Author notes:** These authors contributed equally to this work (Maya Juman, Jeffrey B. Doty).

## Abstract

Borealpox virus (BRPV; formerly Alaskapox) is an orthopoxvirus that has caused seven reported human infections in Alaska since 2015, including a fatal case in 2023. The natural reservoir of BRPV is unknown, although previous investigations have raised the possibility of wild small mammals transmitting the virus to humans, either through direct contact or via domestic cats and dogs. To understand which species may be involved in the maintenance and/or spillover of BRPV in Alaska, we trapped and sampled wild small mammals (including voles, shrews, and squirrels) in 2021 and 2024 near reported human case locations in Fairbanks and the Kenai Peninsula, respectively. We found evidence of previous exposure to orthopoxviruses in five species (including the House Mouse, *Mus musculus*) and detected BRPV DNA as well as viable virus in Northern Red-backed Voles (*Clethrionomys rutilus*). Further, screening of tissues from historical museum specimens revealed BRPV DNA in *C. rutilus* specimens collected in Denali National Park and Preserve in 1998 and 1999, 17 years before the first reported human case of BRPV. Phylogenomic analysis of all human and animal BRPV isolates strongly supports the hypothesis of local human infections through multiple spillover events. These findings suggest *C. rutilus* as a possible reservoir species for BRPV and indicate that BRPV has been present in Alaskan wild small-mammal populations for at least 25 years. Our study highlights the potential of museum collections to elucidate the temporal, spatial, and host ranges of emerging pathogens. Further museum- and field-based sampling will clarify the true geographic range of BRPV, which is closely related to Old World orthopoxviruses and may be circulating beyond North America.

## Introduction

The genus *Orthopoxvirus* (family *Poxviridae*) contains ten recognized species of double-stranded DNA viruses that cause disease in a wide range of mammalian hosts, including humans (Bonwitt et al. 2022). Notably, orthopoxviruses (OPXVs) include variola virus (VARV), which causes smallpox, and monkeypox virus (MPXV), the causative agent of monkeypox, which has recently caused widespread human outbreaks with sustained human-to-human transmission (Bonwitt et al. 2022; WHO 2022). OPXVs may be emerging at higher rates due to increased host mixing and human-wildlife contact driven by climate and land-use change (Thomassen et al. 2013), as well as heightened human susceptibility following the cessation of smallpox vaccination after its global eradication in 1980 (Diaz 2021).

The most recently described OPXV is borealpox virus (BRPV), formerly known as Alaskapox virus (Gigante et al. 2019). BRPV was first detected after a patient presented with a lesion near Fairbanks, Alaska, USA in July 2015 (Springer et al. 2017). Over the following eight years, five additional, epidemiologically unrelated cases were reported in Fairbanks, all involving self-limiting illness (Mooring et al. 2020; Mooring et al. 2021; Mooring et al. 2025). However, in September 2023, the first fatal case was reported in an elderly, immunocompromised patient on the Kenai Peninsula in Southcentral Alaska (∼320 miles straight-line distance SSE of Fairbanks), expanding the known geographic range of this virus and demonstrating its potential for causing fatal outcomes, particularly in severely immunocompromised individuals (Rogers et al. 2025). Furthermore, the viral sequence recovered from this patient was distinct from sequences isolated from prior cases in the Fairbanks area (Rogers et al. 2025), suggesting genetic variation across space as is the case in OPXVs with wide host ranges such as cowpox virus (CPXV) and zoonotic MPXV (Reynolds et al. 2018). Unraveling the geographic origins and range, host species, and transmission pathways of BRPV is important for understanding and mitigating this viral pathogen.

In the initial 2015 investigation of the first BRPV case, inadvertent viral importation from outside of Alaska could not be excluded (Springer et al. 2017). However, the subsequent cases in the Fairbanks area, the patients’ lack of out-of-state travel, and the greater genetic diversity seen in the Kenai isolate strongly indicate that the virus is circulating in at least one non-human reservoir in Alaska. This is particularly noteworthy given the phylogenetic relationship between BRPV and other OPXVs. The genus *Orthopoxvirus* contains two reciprocally monophyletic clades: “Old World” and “New World” OPXVs, which are thought to have diverged from a common ancestor approximately 42,000 years ago (Babkin et al. 2022). Despite being detected only in Alaska, BRPV isolates share a most recent common ancestor with Old World OPXVs and constitute a reciprocally monophyletic sister clade with an estimated divergence date of approximately 19,000 years ago (Babkin et al. 2022). The geographic range of BRPV remains unknown, and it may be circulating beyond Alaska and North America.

The natural reservoir (sensu Viana et al. 2014) of BRPV is also unknown, but evidence from cases, wildlife screening, and the genomic diversity within BRPV suggests small mammals play a role (Mooring et al. 2025). All known patients lived in low population-density, forested areas and reported contact with domestic animals (i.e., dogs and cats), many of which were known to hunt small mammals such as rodents and shrews (i.e., pet dogs and cats that purportedly hunted small mammals) (Springer et al. 2017; Mooring et al. 2020; Mooring et al. 2021; Mooring et al. 2025; Rogers et al. 2025). Furthermore, all reported cases occurred in the late summer to early fall, when outdoor recreational and subsistence activities (e.g., hunting and foraging) increase across Alaska; three out of the seven patients reported berry picking prior to symptom onset. These activities increase opportunities for direct wildlife contact or indirect exposure via fomites, both established pathways for OPXV transmission (Bonwitt et al. 2022). In 2015, samples from small mammals collected near the first patient’s home were screened for OPXV DNA by PCR (serological testing was not conducted); all samples were negative (Springer et al. 2017). During a 2020 investigation following a reported case in the Fairbanks area, 176 small mammals were captured and sampled. Anti-OPXV IgG antibodies were detected in serum samples from 28 of 147 Northern Red-backed Voles (*Clethrionomys rutilus*; 19.0%), one of four Northern Flying Squirrels (*Glaucomys sabrinus*; 25.0%), and one of three American Red Squirrels (*Tamiasciurus hudsonicus*; 33.3%) (Mooring et al. 2025). Viral DNA was amplified from tissue samples of 12 of 149 Northern Red-backed Voles (8.1%) and one of 14 Masked Shrews (*Sorex cinereus*; 7.1%). Among PCR-positive samples, viable BRPV was detected in one Northern Red-backed Vole and one Masked Shrew.

The emergence of BRPV in North America raises pressing questions about its origins and actual geographic distribution, which additional field-based sampling can help address. A complementary approach involves the screening of historical specimens in natural history museums. These collections are valuable but largely underutilized resources for pathogen reservoir identification, with the potential to expand the known spatial, taxonomic, and temporal distributions of viruses of concern (Yates et al. 2002; Colella et al. 2021). In this study, we aimed to identify the reservoir host(s) for BRPV through both field- and museum-based screening. First, we sampled small mammals in and around the properties of human cases and other non-epidemiologically linked locations near Fairbanks and Kenai, Alaska. Here, we sought to expand the taxonomic and geographic scope of previous sampling efforts (Springer et al. 2017; Mooring et al. 2025) by screening a larger sample of small mammals, following similar field studies of MPXV and Akhmeta virus, another recently described OPXV (Doty et al. 2017; Doty et al. 2019). To broaden the temporal scope, we also screened tissue samples from historical specimens housed at the University of Alaska Museum (UAM), targeting species based on preliminary results from field-based sampling (Mooring et al. 2025).

## Results

### Small mammals trapped in 2021 and 2024

In September 2021, 202 wild small mammals representing at least six species were trapped and sampled in the Fairbanks and North Pole regions of Interior Alaska (Table 1). The majority were Northern Red-backed Voles (*n* = 147), followed by shrews (*Sorex* spp.; 21), American Red Squirrels (15), Northern Flying Squirrels (12), voles (*Microtus* sp.; 6), and a single Woodchuck (*Marmota monax*). Additionally, a series of domestic/commensal rats (*Rattus* sp.; 7) obtained by the Alaska Department of Fish and Game (and subsequently deposited in the UAM’s Mammal Collection) from the Fairbanks area were also sampled. In August 2024, 135 wild small mammals of at least six species were trapped and sampled in the Kenai Peninsula area of Alaska (Table 1). Similarly, the majority were Northern Red-backed Voles (83), followed by shrews (37), American Red Squirrels (11), Northern Bog Lemmings (*Synaptomys borealis*; 2), one House Mouse (*Mus musculus*), and one Ermine (*Mustela erminea*).

**Table 1.** Results of serological assays, PCR assays, and viral culture attempts on samples taken from wild small mammals trapped in 2021 and 2024. A breakdown of results by trap site is included in Supplementary Data S1.

| Common name | Species | Sample size | ELISA positives (%) | PCR positives (%) | Viable virus (%) |
| --- | --- | --- | --- | --- | --- |
| Fairbanks and North Pole, Alaska; September 2021 |  |  |  |  |  |
| Northern Red-backed Vole | *Clethrionomys rutilus* | 147 | 20 (13.6) | 7 (4.8) | 0 |
| Shrew | *Sorex* spp. | 21 | 0 | 0 | 0 |
| American Red Squirrel | *Tamiasciurus hudsonicus* | 15 | 5 (33.3) | 0 | 0 |
| Northern Flying Squirrel | *Glaucomys sabrinus* | 12 | 7 (58.3) | 0 | 0 |
| Rat | *Rattus* sp. | 7 | 0 | 0 | 0 |
| Vole | *Microtus* sp. | 6 | 0 | 0 | 0 |
| Woodchuck | *Marmota monax* | 1 | 0 | 0 | 0 |
| Totals |  | 209 | 32 (15.3) | 7 (3.3) | 0 |
| Kenai Peninsula, Alaska; August 2024 |  |  |  |  |  |
| Northern Red-backed Vole | *Clethrionomys rutilus* | 83 | 7 (8.4) | 1 (1.2) | 1 (1.2) |
| Shrew | *Sorex* spp. | 37 | 1 (2.7) | 0 | 0 |
| American Red Squirrel | *Tamiasciurus hudsonicus* | 11 | 1 (9.1) | 0 | 0 |
| Northern Bog Lemming | *Synaptomys borealis* | 2 | 0 | 0 | 0 |
| House Mouse | *Mus musculus* | 1 | 1 (100.0) | 0 | 0 |
| Ermine | *Mustela erminea* | 1 | 0 | 0 | 0 |
| Totals |  | 135 | 10 (7.4) | 1 (0.7) | 1 (0.7) |

In 2021, 20 Northern Red-backed Voles (13.6%) were IgG-positive for OPXV antibodies by ELISA, as were five American Red Squirrels (33.3%) and seven Northern Flying Squirrels (58.3%). However, OPXV DNA was only detected in seven Northern Red-backed Voles (4.8%). In 2024, seven Northern Red-backed Voles (8.4%) were IgG-positive for OPXV antibodies by ELISA, as were one shrew (2.7%), one American Red Squirrel (9.1%), and one House Mouse (100%). Once again, the only PCR-positive animal was a Northern Red-backed Vole (1.2% of 83 *C. rutilus*). Viable virus was isolated from one PCR-positive Northern Red-backed Vole sample collected in 2024. Sequencing was conducted on this viral isolate and one PCR-positive Northern Red-backed Vole sample from 2021. None of the PCR-positive animals from either sampling year had visible lesions suggestive of a poxvirus infection.

### Historical voucher specimen investigations

We screened a total of 285 frozen tissue samples from vole, shrew, and squirrel specimens housed at UAM for OPXV DNA. Four of the 201 samples of Northern Red-backed Voles were positive for OPXV DNA (Table 2). Two of these four PCR-positive samples were taken from the same vole specimen (UAM 51567), and both yielded viable virus (Table 3). All three specimens (UAM 51528, UAM 51567, UAM 73062) were collected at Rock Creek Site in Denali National Park and Preserve, Alaska, in July 1998, August 1998, and August 1999, respectively (Fig. 1). A frozen liver sample was positive from each specimen, as well as a frozen mixed-tissue sample from UAM 51567. Frozen spleen and mixed-tissue samples from UAM 73062 were also screened, and neither was positive for OPXV DNA.

**Table 2.** Results of PCR and viral culture attempts on samples from museum voucher specimens. Collection date, locality, and tissue type of each sample are included in Supplementary Data S1.

| Common name | Species | Samples | PCR positives (%) | Viable virus (%) |
| --- | --- | --- | --- | --- |
| Northern Red-backed Vole | *Clethrionomys rutilus* | 201 | **4 (2.0)** | **2 (1.0)** |
| Tundra Vole | *Alexandromys oeconomus* | 22 | 0 | 0 |
| Insular Vole | *Microtus abbreviatus* | 6 | 0 | 0 |
| Vole | *Microtus* sp. | 1 | 0 | 0 |
| Masked Shrew | *Sorex cinereus* | 43 | 0 | 0 |
| Tundra Shrew | *Sorex tundrensis* | 8 | 0 | 0 |
| Montane Shrew | *Sorex monticolus* | 3 | 0 | 0 |
| American Red Squirrel | *Tamiasciurus hudsonicus* | 1 | 0 | 0 |
| Totals |  | 285 | **4 (1.4)** | **2 (0.7)** |

**Table 3.** Tissue type, collection date, and locality of OPXV PCR-positive Northern Red-backed Vole (*Clethrionomys rutilus*) museum voucher specimens.

| Catalog number | Tissue type | Viable virus isolated? | Locality | Coordinates | Collection date | Sex |
| --- | --- | --- | --- | --- | --- | --- |
| UAM 51528 | Liver | No | Rock Creek Site, Denali National Park, Alaska | 63.729636, -148.982167 | 21 July 1998 | Female |
| UAM 51567 | Liver | Yes | Rock Creek Site, Denali National Park, Alaska | 63.727261, -148.980936 | 20 August 1998 | Male |
|  | Mixed tissue | Yes |  |  |  |  |
| UAM 73062 | Liver | No | Rock Creek Site, Denali National Park, Alaska | 63.730683, -148.984308 | 31 August 1999 | Male |

**Fig. 1.**
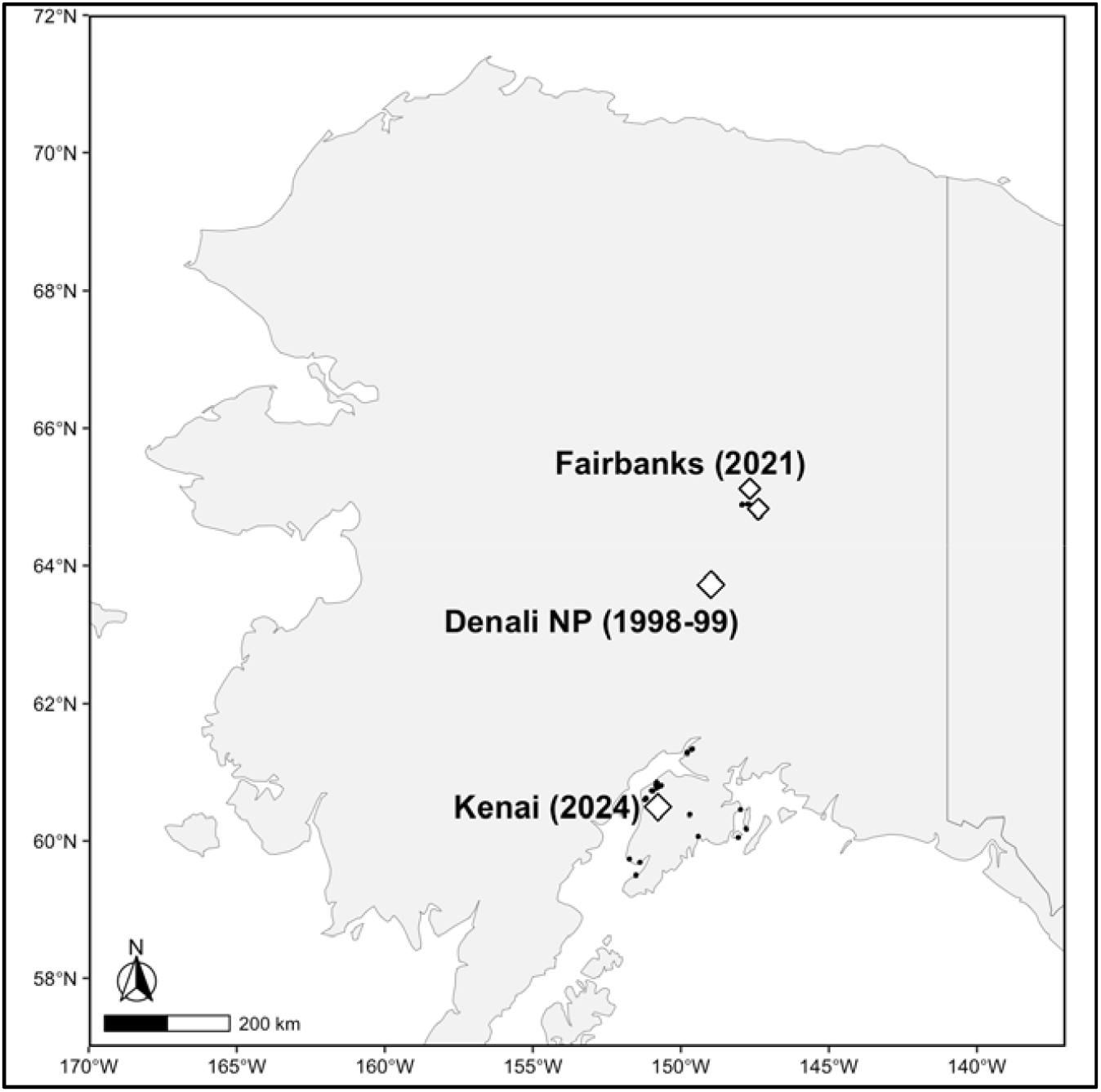
Localities of museum specimens sampled in our study. Diamonds indicate PCR-positive specimens and smaller black dots indicate all other sampling sites.

### Phylogenetic analysis of borealpox virus whole-genome sequences

A whole-genome maximum-likelihood tree revealed that isolates from Fairbanks human cases (2015-2023) form a well-supported clade with small mammal specimens from Fairbanks *C. rutilus* (2020, 2021) and *S. cinereus* (2020) sampled in this and an earlier study (Mooring et al. 2025). There was also strong support for a separate clade containing isolates from the 2023 Kenai human case and *C. rutilus* sampled from Kenai in 2024. The BRPV isolate from the *C. rutilus* specimen collected in Denali National Park and Preserve in 1998 (UAM 51567) fell within the Fairbanks clade, clustering with human cases from 2022 and 2023 (Fig. 2). Notably, both Bayesian and maximum-likelihood analyses did not support monophyly of human-derived sequences, suggesting multiple separate spillover events.

**Fig. 2.**
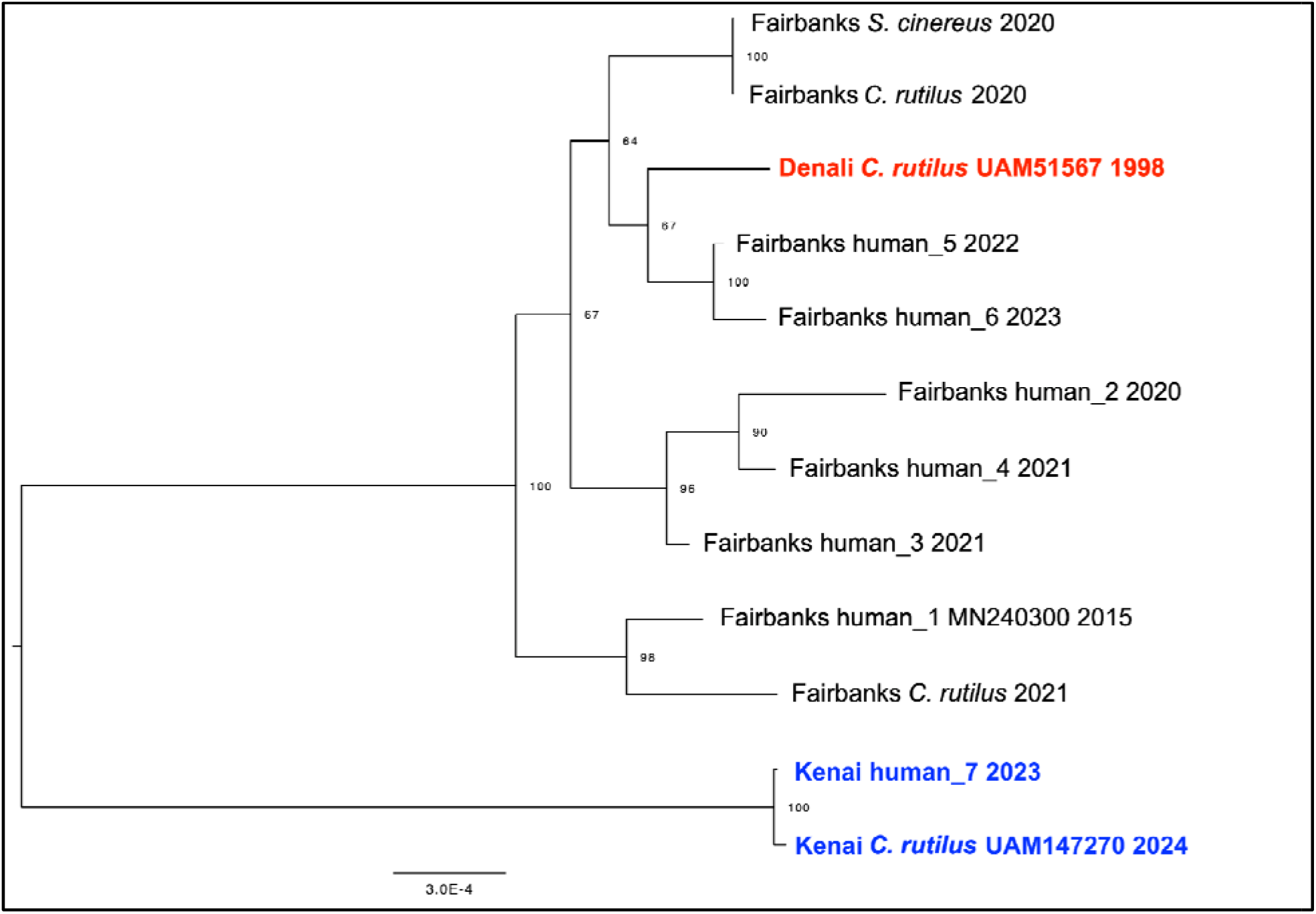
Maximum-likelihood tree based on whole-genome sequences from BRPV isolates, constructed using a GTR+F model in IQ-TREE (Nguyen et al. 2015). Blue labels indicate human and vole isolates from the Kenai Peninsula, while the red label indicates the isolate from UAM 51567. The vole sequence from Fairbanks 2021 was sequenced directly from DNA extracted from tissue. Bootstrap support values are shown at each node.

## Discussion

Borealpox virus (BRPV) is a recently identified OPXV that has repeatedly spilled over into human populations in Alaska since first reported in 2015. To better understand the zoonotic reservoir of BRPV, we trapped and sampled wild small mammals in areas of known BRPV spillover in Alaska—the Interior Region’s Fairbanks region in September 2021 and the Kenai Peninsula in 2024—and screened tissue and blood samples for OPXV DNA and OPXV antibodies, respectively. We also screened tissue samples associated with historical museum specimens of suspected host species (collected 1991-2019 and housed at UAM) for OPXV DNA.

As in earlier field-based surveys (Mooring et al. 2025), most seropositive animals in our study were Northern Red-backed Voles (*n* = 27), with 13.6% seropositivity and 8.4% seropositivity for OPXV antibodies in Fairbanks (2021) and Kenai (2024), respectively. American Red Squirrels, Northern Flying Squirrels, an unspecified shrew, and a House Mouse were also seropositive (Table 1). The IgG ELISA used in this study is not BRPV-specific, so these results may reflect past infection with other OPXVs, which could potentially circulate undetected in these small-mammal populations; however, no other OPXVs have been identified to date in Alaska. Other OPXVs thought to circulate in North American small mammals have not been detected in Alaska, including raccoonpox, volepox, and skunkpox viruses (Alexander et al. 1972; Regnery 1987; Emerson et al. 2009). Consequently, the detection of OPXV antibodies in a House Mouse (*M. musculus*) is notable; should this reflect a prior BRPV infection, it suggests the potential for mouse-human transmission risk in domestic settings, which may be higher than the risk of transmission from other small mammals that are less likely to be found in human dwellings. The distribution and prevalence of *M. musculus* in Alaska is very poorly understood, but museum voucher specimens have been collected from or near Fairbanks, Fort Yukon, Anchorage, the Kenai Peninsula, Kodiak Island, St. Paul Islands, Unalaska Island, Kiska Island, and multiple localities in Southeast Alaska (MacDonald and Cook 2009). Predictive modeling based on viral genomic features has implicated the genus *Mus* as potentially suitable borealpox hosts (Tseng et al. 2025). Expanded serological and molecular testing of mice in and around the homes of BRPV patients, and throughout Alaska, will be critical for investigating this hypothesis.

Only Northern Red-backed Voles tested positive for OPXV DNA (seven in 2021 and one in 2024), two of which were confirmed as BRPV by DNA sequencing (Figure 2.) Viable virus was isolated from the positive sample collected in 2024. This supports prior evidence that the Northern Red-backed Vole, which is also commonly found in and around human dwellings, is involved in the maintenance and circulation of BRPV in wildlife populations in Alaska (Mooring et al. 2025). To further investigate the reservoir of BRPV, future research should include longitudinal study of potential host species, confirmation of long-term viral maintenance in these populations, and expanded screening of species beyond Northern Red-backed Voles. Our study is limited by small sample sizes of most species and a heavy bias toward *C. rutilus* based on the availability of samples to screen and previous evidence of BRPV circulation in this species.

Other species—including squirrels, a mouse, and a shrew—were seropositive in our study, and the screening of larger sample sizes of these species may reveal evidence of PCR positivity and viable virus. A recent study suggests squirrels as possible reservoirs of MPXV, a related OPXV (Riutord-Fe et al. 2026), and future work on BRPV should continue to investigate the possibility of other small mammal species playing a role in its maintenance and transmission.

While future field studies of BRPV dynamics in wild mammals will be critical for elucidating the reservoir species, looking for evidence of past BRPV circulation in Alaska can complement contemporary data and further our understanding of the ecology of this virus. To do this, we screened 285 tissue samples from historical museum specimens of voles, shrews, and squirrels for OPXV DNA, and found four positive samples—all from Northern Red-backed Voles (Table 2). DNA sequencing confirmed the presence of BRPV DNA in one of these samples (Figure 2.). Positive samples were collected from Denali National Park and Preserve in 1998 and 1999, over 15 years before the first reported case of human BRPV infection (Table 3). Denali National Park and Preserve is roughly equidistant from the known spillover locations of BRPV (Fairbanks and Kenai Peninsula) (Fig. 1). The detection of BRPV DNA in samples from the 1990s and in a different site from known cases confirms the wider circulation of BRPV in Alaskan small-mammal populations over a longer period of time than previously known. Phylogenetically, the isolate from this museum specimen fell within a clade of contemporary human and small-mammal isolates from the Fairbanks area (Fig. 2). Another separate, reciprocally monophyletic clade contains isolates from the Kenai Peninsula human case in 2023 and *C. rutilus* sampled from case patient’s residence in 2024, suggesting that the Kenai patient was likely infected locally (Fig. 2). Our findings support the hypothesis of a long-term, established wildlife reservoir with occasional spillover into human populations and potentially other mammal populations. Initial investigations of human cases could not exclude the possibility of recent BRPV introduction into Alaska (Springer et al. 2017). However, our study suggests that BRPV has been present in Alaska for at least 25 years (and likely much longer) with multiple separate spillover events since 2015, though the drivers of these events remain unknown. It is possible that previous spillover events went undetected due to mild, self-limiting illness or symptoms being mistaken for other rashes (Mooring et al. 2020; Mooring et al. 2021). Alternatively, spillover may have become more likely in the past decade due to the effects of climate and land-use change on the distributions and cross-species interactions of small mammals (Baltensperger et al. 2024), as is the case for MPXV (Thomassen et al. 2013), and/or waning population-level immunity against OPXVs following the cessation of smallpox vaccination (Diaz 2021).

The Northern Red-backed Vole is found in Alaska, Canada, northern Russia, and Scandinavia (Linzey et al. 2020). If this species is the primary reservoir of BRPV, our findings suggest that BRPV may be circulating beyond North America, in *C. rutilus* as well as in related species that hybridize with *C. rutilus* in contact zones (Runck et al. 2009; Wiens and Colella 2025). A broader awareness of BRPV symptoms and diagnostics will be critical to identify other spillover events in Alaska and the broader arctic region. Further molecular and serological surveillance of wild mammal populations in Alaska, as well as longitudinal studies of suspected host species like Northern Red-backed Voles, will be vital to identifying the drivers of the apparently sudden spillover of BRPV into human populations.

When federal public health agencies partner with non-governmental organizations such as natural history museums and academic institutions, investigations are strengthened by leveraging the resources, capabilities, and expertise of all partners. Our study illustrates the value of museum specimens and their unique potential to elucidate the origins, reservoirs, and dynamics of emerging viruses. As demonstrated here, disease surveillance in historical museum collections can expand our understanding of the temporal, spatial, or host taxonomic range of newly discovered pathogens (Juman et al. 2025; Cronin et al. 2025). This work can clarify whether a virus was recently introduced to a given region or whether it has been endemic for decades and present in additional nearby areas, as is the case for BRPV in Alaska. The continued collection and preservation of voucher specimens will allow future researchers to retrospectively understand pathogen emergence in the context of unprecedented global change.

## Methods

### Small-mammal trapping and specimen collection

Small mammals were trapped at nine sites around the Fairbanks and North Pole regions of Alaska during September 6-14, 2021 (Fig. 1). Sites included homes of the two human cases reported in 2021 and locations within seven miles of the case’s home where permission was received from landowners. Other potential exposure locations such as homes of family members were also included. All sites were in mixed evergreen and deciduous forests, either on public land (*n* = 2) or low-density residential areas (*n* = 7).

Following the identification of the BRPV case on the Kenai Peninsula in late 2023, small mammals were also trapped at three sites in the Kenai/Soldotna/Sterling area during August 15-22, 2024 (Fig. 1). One site was on public land north of the Kenai Municipal Airport and was partially boggy while the other two sites were in low-density residential areas with mixed evergreen and deciduous forests, including the home of the case patient.

Trapping was primarily conducted using standard (7.6 x 8.9 x 22.9 cm) Sherman live traps (H.B. Sherman Traps, Inc., Tallahassee, FL, USA), which were deployed for a total of 1384 trap-nights (i.e., the sum of the number of traps placed each night over the course of trapping) in 2021 and 1825 trap-nights in 2024. Tomahawk live traps (Model 102, Tomahawk Live Trap, Hazelhurst, WI, USA) were used for larger mammals such as squirrels, and pitfall traps were deployed to catch shrews. The Tomahawk and Sherman traps were baited with a mixture of peanut butter and oats, with carrots included in Fairbanks due to colder temperatures. Trapped animals were anesthetized with isoflurane then humanely euthanized and examined prior to examination for skin lesions and collection of blood and tissue samples (liver, skin, pooled heart and lung, and pooled kidney and spleen; lesion material if observed). All trapping and animal handling followed protocols approved by the CDC and UAF’s Institutional Animal Care and Use Committees (CDC: 3183DOTMULX, 3400DOTMULX; UAF: 152295). Tissue specimens were also obtained from roadkill animals collected by the UAM researchers and from animals collected by the Alaska Department of Fish and Game. All voucher specimens and their associated tissues have been deposited and catalogued in UAM’s Mammal Collection (Supplementary Data S1).

### Museum specimen sampling

We screened tissue samples from specimens of voles, shrews, and squirrels archived at UAM. The following species were selected for screening based on the availability of tissue samples, as well as preliminary results about seropositivity and molecular BRPV detection in wild small mammals: Northern Red-backed Voles (*n* = 201 samples), Tundra Voles (*Alexandromys oeconomus*; 22), Singing Voles (*Microtus miurus*; 6), a vole of undetermined species (*Microtus* sp.; 1), Masked Shrews (43), Tundra Shrews (*Sorex tundrensis*; 8), Montane Shrew (*Sorex monticolus*; 3), and an American Red Squirrel (1). The specimens were collected at various locations around Fairbanks, Kenai, and Denali National Park and Preserve from 1991-2019 (Fig. 1; Supplementary Data S1).

Historical samples archived at UAM were flash-frozen without a buffer and housed in liquid nitrogen-cooled cryovats that maintain vapor-phase nitrogen at -170°C. The work surface and tools were cleaned with 10% bleach. The box containing the appropriate tissues was removed from the cryovat freezer and placed on an insulated cold-block while outside the cryovat. Using a separate set of cleaned instruments for each tube, a small subsample (roughly 0.1 gram) of tissue was removed from each cryotube and transferred into appropriately labeled tubes. Between sampling, instruments were cleaned in 10% bleach solution and then rinsed with water and dried. After subsampling, the original samples were returned to the cryovat freezer, and the subsamples were placed in a labeled box on a second insulated cold-block. Once all subsampling was completed, the box of subsamples was stored in an ultra-low freezer until shipment. Samples were shipped on dry ice.

### Laboratory Diagnostics

Tissue samples were homogenized in 500μl of PBS with 250μl of 1mm zirconia silica beads (Biospec) and a Mini BeadBeater 24 following the manufacturer’s recommendations. 100μl of tissue homogenate was used for DNA extraction with a MagMAX deep-well magnetic processor (ThermoFisher Scientific, https://www.thermofisher.com) using the MagMAX DNA Multi-Sample Ultra kit. DNA samples were assessed for the presence of OPXV with the CDC OPXV generic real-time PCR assay (Li et al. 2006).

A modified anti-OPXV IgG enzyme-linked immunosorbent assay (ELISA) was conducted on sera obtained from cardiac punctures or on dried blood spots collected on Nobuto filter paper (Advantec, San Diego, CA) as previously described (Hutson et al. 2009, Doty et al. 2017). Samples were considered positive if the OD value passed the cut-off value in at least two consecutive dilutions (1:100 and 1:200).

### Viral isolation and DNA sequencing

Following PCR, samples with OPXV DNA amplification were added to cell culture for virus isolation. Remaining tissue homogenate from the DNA extraction process was added to BSC-40 cell monolayers (African Green Monkey *Chlorocebus sabaeus* kidney cell line) in T-25 cell culture flasks and incubated at 37°C with 5% CO2 (Hutson et al. 2013). DNA was then extracted from propagated virus and sheared to target 500 bp fragments on a Covaris S220 instrument (Covaris, Woburn, Massachusetts, USA). The Swift Accel-NGS 2S DNA Library Kit was used for library preparation with dual indexing following the manufacturer’s instructions (Swift Biosciences, product no longer available). Libraries were visualized with an Agilent 2200 Tape Station (Agilent Technologies, Santa Clara, California, USA), followed by sequencing on an Illumina MiSeq machine with the MiSeq Reagent v3 600 cycle kit (Illumina, San Diego, California, USA).

### Genome Assembly

Reads were trimmed to a minimum length of 50 bp and minimum quality 20 with FaQCs v1.34 (https://github.com/LANL-Bioinformatics/FaQCs), and three nucleotides were removed from each end. The 2015 BRPV reference genome (GenBank accession MN240300.1) was used for mapping, with bwa mem v0.7.17 (https://github.com/lh3/bwa). Mapped reads were then extracted with samtools v.1.9 (https://github.com/samtools/samtools) and *de novo* assembled with SPAdes v3.13.0 (https://github.com/ablab/spades), using the --careful flag and --cov-cutoff off. Contigs with coverage >10 were manually assembled in Geneious Prime 2023.0.4 (Dotmatics, Boston, Massachusetts, USA) after mapping. Draft genomes were then edited in repeat regions at approximate positions 150, 163, 174, and 200 kb to match repeat lengths from the 2015 BRPV genome, and only the left ITR and 500 bp of the right ITR were retained. Trimmed reads were mapped back to the draft genomes, and a polished genome was generated with ivar v0.1, using the conditions above except requiring a minimum read depth of 10. Finally, to produce a final genome, ITRs were copied from the left to the right end. Annotations from the reference genome were transferred using Liftoff (https://github.com/agshumate/Liftoff).

### Phylogenetic Analysis

Genome alignments were performed with MAFFT v7.490 in Geneious Prime 2023.0.4 using FFT-NS-i x2 and final alignments are provided as supplementary files (Supplementary Data S2). Bayesian phylogenetic trees were generated in BEAST v2.7.7 (Bouckaert et al. 2019) in two runs with the following parameters based on Gigante et al. (2019): GTR+I+G nucleotide substitution model (4 gamma categories, 35% invariant), relaxed lognormal clock (exponential distribution of ucldStdev prior with mean = 0.333), and Yule model prior until all parameters exhibited ESS > 200 after 10% burn-in, visualized in Tracer. Default parameters were used unless specified. Run log and tree files were combined using LogCombiner after 10% burn-in. A maximum clade credibility tree was estimated in TreeAnnotator based on sampling frequency of 1000 and 10% burn-in. We also estimated a maximum-likelihood tree on the whole genome using IQ-TREE (Nguyen et al. 2015), with branch supports obtained using the ultrafast bootstrap method and 1000 replicates. The nucleotide substitution model for the maximum-likelihood tree (GTR+F) was determined using ModelFinder implemented through IQ-TREE (Kalyaanamoorthy et al. 2017). Vaccinia virus (GenBank AY603355) was used as an outgroup.

## Acknowledgements

This research was supported by generous donations to the University of Alaska Museum’s Mammal Collection from the Jay Pritzker Foundation and Nancy Eliason. MMJ was supported by a Gates Cambridge Scholarship enabled by grant OPP1144 from the Bill & Melinda Gates Foundation. CNM, AM, and AC were supported in part by appointment to the Research Participation Program at the Centers for Disease Control and Prevention, administered by the Oak Ridge Institute for Science and Education through an interagency agreement between the U.S. Department of Energy and CDC. MMM was supported by an NSF CAREER Award (2238801). The authors thank Mallory Gulbranson and Kyndall B.P. Hildebrandt at the University of Alaska Museum for access to frozen tissues in their care and assistance with subsampling. The authors also thank Nicholas Fowler from Alaska Department of Fish and Game for his help with accessing sampling sites on the Kenai Peninsula.

## Disclaimer

The findings and conclusions in this report are those of the authors and do not necessarily represent the official position of the Centers for Disease Control and Prevention.

